# An insula-driven network computes decision uncertainty and promotes abstinence in chronic cocaine users

**DOI:** 10.1101/2020.04.08.031757

**Authors:** Ju-Chi Yu, Vincenzo G. Fiore, Richard W. Briggs, Jacquelyn Braud, Katya Rubia, Bryon Adinoff, Xiaosi Gu

## Abstract

The anterior insular cortex (AIC) and its interconnected brain regions have been associated with both addiction and decision-making under uncertainty. However, the causal interactions in this uncertainty-encoding neurocircuitry and how these neural dynamics impact relapse remain elusive. Here, we used model-based fMRI to measure choice uncertainty in a motor decision task in 61 individuals with cocaine use disorder (CUD) and 25 healthy controls. CUD participants were assessed before discharge from a residential treatment program and followed for up to 24 weeks. We found that choice uncertainty was tracked by the AIC, dorsal anterior cingulate cortex (dACC), and ventral striatum (VS), across participants. Stronger activations in these regions measured pre-discharge predicted longer abstinence after discharge in individuals with CUD. Dynamic causal modelling revealed an AIC-to-dACC directed connectivity modulated by uncertainty in controls, but a dACC-to-AIC connectivity in CUD participants. This reversal was mostly driven by early-relapsers (<30 days). Furthermore, CUD individuals who displayed a stronger AIC-to-dACC excitatory connection during uncertainty encoding remained abstinent for longer periods. These findings reveal a critical role of an AIC-driven, uncertainty-encoding neurocircuitry in protecting against relapse and promoting abstinence.

## 1. Introduction

Despite the high public health burden of substance use disorders (SUDs), existing treatments are often ineffective. Thus, relapse - the reinstatement of compulsive drug-seeking behaviors after drug abstinence - remains common (Kelley, 2004). Risky decision-making has been extensively studied as a behavioral phenotype associated with increased relapse vulnerability (McLellan *et al*., 2000; Gowin *et al*., 2013; Koob & Volkow, 2016; Verdejo-Garcia *et al*., 2018). These studies typically require the subject to choose between ‘safe’ and ‘risky’ options (Lane & Cherek, 2000; Fishbein *et al*., 2005; Xiao *et al*., 2012; Orsini *et al*., 2017), and have found that both substance-dependent humans and non-human animals are more likely to choose risky options over safe options in comparison with non-addicted controls (Ersche *et al*., 2005; Mitchell *et al*., 2014). Neurally, it has also be shown that risky choices elicit stronger responses in subcortical regions (e.g. striatum) but weaker prefrontal activities in methamphetamine-dependent humans compared to healthy controls (Ersche *et al*., 2011; Kohno *et al*., 2014).

A key computational process involved in decision-making is the estimation of action-outcome probabilities (De Martino *et al*., 2013; Payzan-LeNestour *et al*., 2013; Meyniel *et al*., 2015). That is, an individual must be able to accurately represent how likely her chosen action is to lead to a desirable result: we defined this quantity as choice uncertainty, following previous literature (Meyniel *et al*., 2015; Adler & Ma, 2018; Atiya *et al*., 2020). Previous studies have shown the involvement of several brain regions in the estimation of such probabilities, including the anterior insular cortex (AIC) (Clark *et al*., 2008; Preuschoff *et al*., 2008), dorsal anterior cingulate cortex (dACC) (Rushworth & Behrens, 2008; Stolyarova *et al*., 2019), and ventral striatum (VS) (Berns *et al*., 2001). For instance, Preuschoff and colleagues used a gambling task where the probability of stimulus-reward associations changed over time. They found that in healthy controls, the AIC not only tracked this probability per se (termed “risk” in the paper) but also errors in such risk prediction (Preuschoff *et al*., 2008). The dACC, a brain region closely connected and often co-activated with the AIC, has been consistently implicated in decision making arbitrations based on conflict monitoring and the associated uncertainty estimations (Rushworth & Behrens, 2008; Stolyarova *et al*., 2019) across tasks such as Stroop, flanker or Simon tasks (Venkatraman *et al*., 2009; Shenhav *et al*., 2013).

The same AIC-related circuitry has also been commonly associated with SUD and addiction-related phenotypes (Goldstein *et al*., 2009; Naqvi & Bechara, 2009; Koob & Volkow, 2016), with the VS particularly implicated in association in cocaine use disorder (CUD) and risk-preference behavioral traits (Ersche *et al*., 2011; Mitchell *et al*., 2014). For example, McHugh and colleagues used resting-state fMRI (rsfMRI) and found reduced insula-striatum coupling in individuals with CUD and especially those who relapsed early (<30 days) (McHugh *et al*., 2013) but enhanced rsfMRI connectivity in the AIC-dACC circuitry in those who managed to stay abstinent (McHugh *et al*., 2016). It has been hypothesized that the insula might drive a network of brain regions in integrating interoceptive states and cognitive control under uncertainty and risk in addiction (Naqvi *et al*., 2014), a hypothesis partially supported by animal models (Koob & Volkow, 2016). However, a causal role of the insula in computing uncertainty – particularly in relation to addition relapse - has not been directly examined or established.

Based on this literature, we hypothesized that the AIC might play a crucial role in driving its interconnected regions (e.g. dACC) in encoding choice uncertainty, and that alteration in the neural dynamics of this network would increase the chances of early relapse. We tested these hypotheses by examining 25 healthy controls (HCs) and 61 participants with CUD. All CUD participants were assessed right before discharge from a residential treatment program and followed up for 24 weeks, or until the first day of any use of cocaine or amphetamine. Early relapse was defined *a priori* as return to use within 30 days after discharge (Adinoff *et al*., 2015; McHugh *et al*., 2016). We used a Bayesian model to quantify choice uncertainty in a motor decision-making task (i.e. stop signal task; **Fig. 1**) and dynamic causal modelling (DCM, Friston *et al*., 2003) was applied to model effective connectivity (i.e. directed influence between neuronal populations) subserving uncertainty encoding. The stop signal task has been traditionally used to examine response inhibition and impulsivity (Rubia *et al*., 2003; Schall *et al*., 2017; Verbruggen *et al*., 2019). However, participants also form beliefs representing the probability for a chosen action to be correct (e.g. left-go vs. stop), given an observed sequence of stimuli and the possibility to receive a stop signal after an unknown time interval (cf. Chikazoe *et al*., 2009; Zandbelt & Vink, 2010). In Bayesian terms, newly acquired information, such as the symbolic visual stimuli of the stop signal task, is used to update prior beliefs (Friston, 2010; Friston *et al*., 2016), which can be consolidated or weakened. Thus, each choice selection can be associated with a degree of uncertainty, representing the estimated probability the chosen action might not be correct (Meyniel *et al*., 2015; Adler & Ma, 2018; Atiya *et al*., 2020). A few studies have adopted a similar Bayesian approach to examine prediction error encoding in methamphetamine-dependent individuals using the stop signal task (Harle *et al*., 2015; Harle *et al*., 2016). Here, we will instead examine subject-specific uncertainty encoding and network dynamics to highlight their contribution to CUD relapse, consistent with the computational psychiatry approach to characterize ‘deep’ cognitive processes (Montague *et al*., 2012; Adams *et al*., 2016).

**Fig. 1.**
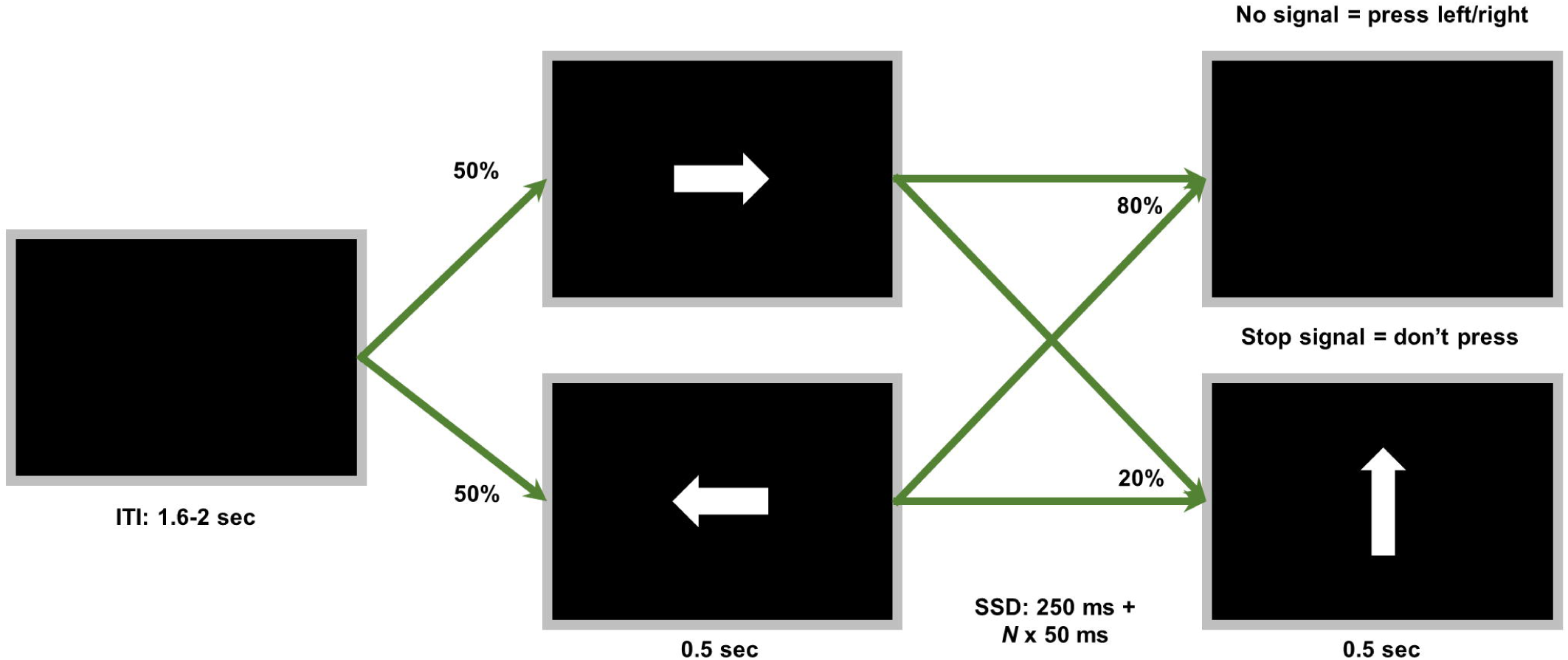
Task details. In this stop signal task, the initial inter-trial interval (ITI) was followed by either a left or a right arrow symbol on the monitor (equal probability for both symbols) for a duration of .5 sec. These initial stimuli were followed, after a variable delay (stop signal delay; SSD), by an upward arrow in 20% of the trials and indicated that the participant should not respond. The VI varied as a function of the behavior of the participants: with a minimum of 250 ms, the VI increased by 50 ms after a successful response inhibition and decreased by 50 ms if any response was produced in a no-response trial.

## 2. Materials and Methods

### 2.1. Participants

We recruited 61 two-to-four weeks abstinent CUD participants, and 25 non-using HC participants with no lifetime history of SUD or other psychiatric disorders (see **Table 1**). All participants provided informed consent prior to the study and received compensation for their participation. The research protocol was reviewed and approved by the Institutional Review Boards of the University of Texas Southwestern Medical Center and the Veterans Administration North Texas Health Care System.

**Table 1.** Demographic data (mean values ± standard deviation)

|  | Healthy controls | CUD participants |
| --- | --- | --- |
| Number of people | 25 | 57 |
| Age (Years) | 39.52 ± 9.29 | 44.04 ± 6.82* |
| Sex |  |  |
| Male | 80 % | 85.96 % |
| Female | 20 % | 14.04 % |
| Ethnic groups |  |  |
| African-American | 56 % | 71.93 % |
| Caucasian | 32 % | 24.56 % |
| Hispanic | 8 % | 0 % |
| Other | 4 % | 3.51 % |
| Education (Years) | 12.56 ± 1.76 | 13.05 ± 1.89 |
| Cocaine Craving Questionnaire (CCQ) | - | 2.49 ± 0.85 |
| Cocaine use (Days) |  |  |
| Life time | - | 3414.22 ± 2215.91 |
| Previous 90 days | - | 65.21 ± 27.09 |
| Money spent for cocaine (\$) | - | |
| Life time | - | 376068.93±<br>410219.51 |
| Previous 90 days | - | 6211.32 ± 5512.29 |

Note: \* indicates $p < .05$ in the Welch's $t$ -test versus healthy control groups.

The residential treatment program for CUD participants lasted two to four weeks. During the first two weeks, CUD participants underwent a comprehensive medical history and physical examination, a general laboratory panel, a urine substance screen, and a guided interview of lifetime substance use history (Kelly *et al*., 2009). The fMRI session took place immediately before CUD participants were discharged from treatment. The fMRI sessions were conducted in the morning at approximately the same time of day for all subjects. Participants were asked to restrict caffeine and nicotine intake to at least one hour before the fMRI session to control for potential acute or withdrawal effects. After discharge, CUD participants were followed for 24 weeks, or until the first day of any use of cocaine or amphetamine, with two appointments per week: one by phone and one in person. A structured interview about substance use and the urine drug screen took place during the in-person clinical appointment. The “days to relapse” measure was defined a priori as either the first day of use of cocaine or amphetamine after discharge (self-report or urine drug screen) or the day of their first missed appointment if participants missed two consecutive appointments and all attempts to contact participant (e.g., phone, mail, collaterals) were unsuccessful..

Sixty-one CUD and 25 healthy participants completed the stop signal task. Four CUD participants were subsequently excluded due to corrupt data archive, which resulted in incomplete data, and one further CUD participant was excluded due to image artifacts. In relapse-related analyses, one CUD participant was also excluded due to the absence of follow-up data. Nine of the CUD participants did not relapse by the end of the follow-up period; their relapse data are here reported as equivalent to 168 days in correlation analyses (i.e. 24 weeks, the duration of the entire follow-up period).

### 2.2. Experimental Design

In the stop signal task (adapted from: Rubia *et al*., 2003), a left or right arrow (randomized) was presented on the screen as the “go signal” in randomly jittered inter-trial intervals of ∼ 1.8s (1.6-2s, **Fig. 1**). The participants were tasked to respond as fast as possible by pressing a left or right button, accordingly, unless the left/right arrow was followed by an upward arrow (in 20% of the trials), which represented the “stop signal”. To ensure equal task difficulty (50% success rate of inhibition) across different participants, the variable interval between the initial left/right arrow and the stop signal (stop signal delay; SSD) changed idiosyncratically by a 50 ms step size, decreasing the interval if the probability of inhibition was below 50%, to make future inhibitions easier for the participant, or increasing the interval if the probability of inhibition was over 50%, to achieve the opposite effect. The task consisted of 362 trials: 288 go and 74 stop trials. The task was presented using the Presentation® software.

### 2.3 Behavioral Analysis

#### 2.3.1. Model-agnostic behavioral analysis

We calculated the standard behavioral indices (Rubia *et al*., 2003) for each group 1) the mean length of the SSD; 2) the mean reaction times (RTs) to go trials and stop failures; 3) The stop-signal RT (SSRT) (Logan *et al*., 2014) for both groups was computed by subtracting the mean length of SSD from the mean RT to go trials.

#### 2.3.2. Computational modeling of behaviors

We used an ideal observer Bayesian model to estimate the trial-by-trial action-outcome distribution of probabilities or *beliefs* associated with the action performed. Given an environment (e.g. visual stimuli) and a set of possible interactions (e.g. press the left-right button), a belief represents the estimated likelihood an action produced in response to a perceived stimulus will result in a known outcome (Friston, 2010; Friston *et al*., 2016), denoted by *p*(*θ*|*a*):

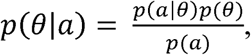

where *a* stands for the action and *θ* for the outcome, i.e. in the present task complying vs not-complying to the task instructions. In this equation, *p*(*θ*) represents the prior probability and *p*(*θ*|*a*) the posterior probability, which is updated depending on each observed outcome, given action *a*. We assumed participants were characterized by normally distributed, subject-specific, beliefs about the distribution of available events/interactions in the task environment *p*(*a*):

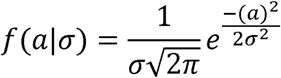

where a different value for the standard deviation (*σ*) was estimated for each participant to determine the pace of belief updates. The model estimated probabilities over two independent dimensions: 1) left/right and 2) go/stop trials, with visual stimuli used as evidence on a trial-by-trial basis to update beliefs about the action to perform (left vs right) and whether to perform an action or not (go vs stop). As a result, the model estimated the subject-specific trial-by-trial probability that each of the available actions (go-left, go-right, stop) would be correct. This estimated quantity determined the confidence (*c*) associated with the chosen action. The complement probability (1-*c*) was used to estimate the probability the chosen action was incorrect: this value was used to represent subject-specific trial-by-trial uncertainty in choice selections. The choice behavior of this task was simulated by the model with a mean prediction accuracy of 90.21±2.59% for HCs with mean *σ*_left/right_ = 0.27±0.05 and *σ*_go/stop_ = 0.44±0.07. The simulation for CUD participants had a mean accuracy of 89.05±5.12% with mean *σ*_left/right_ = 0.29±0.13 and *σ*_go/stop_ = 0.44±0.08.

### 2.4. fMRI data analysis

#### 2.4.1. fMRI acquisition and preprocessing

Anatomical and functional images were collected on a Philips 3T MRI scanner. High-resolution T1-weighted anatomical images were acquired with a 3D magnetization prepared rapid gradient-echo (MPRAGE) sequence with a repetition time (TR) = 8.2 ms, an echo time (TE) = 3.8 ms, a flip angle = 12°, and a 1 × 1 × 1 mm^3^ spatial resolution. Functional images were acquired with a single-shot echo-planar imaging (EPI) sequence with a flip angle of 70°, a TR of 1700 ms, a TE of 25 ms, 36 slices, the field of view (FOV) = 208 × 208 mm^2^, and the voxel size = 3.25 × 3.25 × 3.25 mm^3^. The functional scans were preprocessed using standard statistical parametric mapping (SPM12, Wellcome Department of Imaging Neuroscience; www.fil.ion.ucl.ac.uk/spm/) algorithms, including slice timing correction, co-registration, normalization with resampled voxel size of 2 mm × 2 mm × 2 mm, and smoothing with an 8 mm Gaussian kernel. A temporal high-pass filter of 128 Hz was applied to the fMRI data, and the temporal autocorrelation was modeled using a first-order autoregressive function.

#### 2.4.2. Model-based fMRI data analysis

The first-level general linear model (GLM) included the onsets of button presses as a sensorimotor event (in the case of go trials and unsuccessful stop trials) and the onset of the stop signal (in the case of successful stop trials). Trial-by-trial uncertainty estimated from the Bayesian model was entered as a parametric modulator. The reaction time of each responded trial (or the onset of the stop signal in the successful stop trials) and six motion parameters were included as regressors-of-non-interest in first-level GLMs.

For the ROI analysis, we created masks using the peak coordinates of activation related to uncertainty (*p* < .05, corrected for family-wise error [FWE]) across healthy and CUD groups and examined within their respective CUD groups how these activations were related to relapse. ROI masks were generated as spheres with an 8-mm radius using the MarsBar toolbox (marsbar.sourceforge.net/). Peak coordinates are: bilateral AIC (X = 33/-33, Y = 20, Z = −4), dACC (X = 8, Y = 23, Z = 32), and bilateral VS (left: X = −12, Y = 11, Z = −8; right: X = 13, Y = 13, Z = −8). The mean parameter estimates of GLM were extracted for each participant from each ROI.

#### 2.4.3. Machine learning analysis

To examine how well the uncertainty-related activation can predict participants’ groups, we performed classification tests with leave-one-out support vector machine (SVM). In this SVM, we first extracted, from each participant, uncertainty-related neural signals in all voxels of our chosen ROIs. The classification model was then trained by the extracted ROI activation after leaving one of the participant’s data out, and the trained model was then applied to classify this out-of-sample participant. After repeating this procedure for each participant, a prediction accuracy of the model was obtained. We conducted two types of classifications: 1) early- vs. late-relapse (30 days criterion, as by definition of early-remission in DSM-IV) and 2) HCs vs. CUD participants (**Fig. 2f**).

**Fig. 2.**
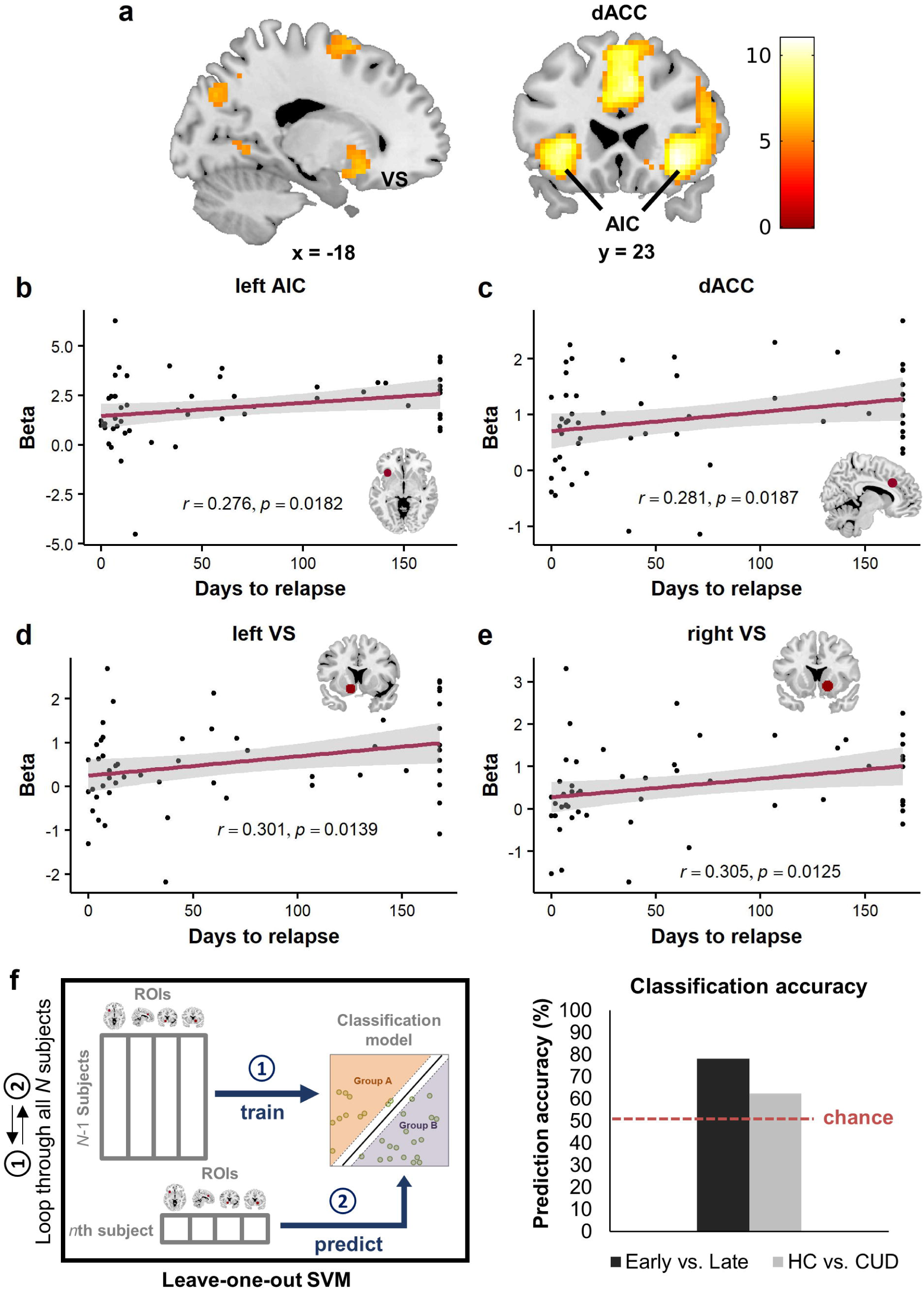
Uncertainty encoding in the stop signal task. **(a)** Across all participants, whole-brain analysis showed uncertainty-related BOLD activation in dorsal anterior cingulate (dACC; extending into supplementary motor area), anterior insula (AIC), and ventral striatum (VS). The image shows BOLD activity FWE corrected *p*<.05, *k*=30. **(b-e)** Beta values extracted from the regions of interest (ROIs) using spherical volumes (r=8mm). This analysis showed significant positive correlations between days to relapse and uncertainty-related activation in **(b)** left AIC (peak coordinates: X = −33, Y = 20, Z = −4), **(c)** right dACC (coordinates: X = 8, Y = 23, Z = 32), **(d)** left VS (coordinates: X = −12, Y = 11, Z = −8), and **(e)** right VS (coordinates: X = 13, Y = 13, Z = −8), suggesting reduced encoding of uncertainty associated with early relapse. **(f)** A leave-one-out support vector machine (SVM) algorithm was conducted to predict participants’ relapse status (early- vs. late-relapse with a 30-day criterion) and diagnostic group membership (healthy controls [HCs] vs. cocaine use disorder [CUD] participants) using their neural encoding of uncertainty. We trained the classifier with the fMRI results extracted from all participants and all voxels of the four ROIs (left AIC, dACC, and bilateral VS). The classification model was first trained after leaving one of the participants out (i.e., with *N*-1 participants’ data). Next, this classification model was applied to predict which group the left-out participant belongs to. After this procedure had been repeated for all participants, a prediction accuracy of this model was obtained: early- versus late-relapse was predicted with 78.18%.

#### 2.4.4. Dynamic causal modelling

We used SPM12 to estimate the effective connectivity among the three key ROIs of the AIC, dACC and VS. We relied on the same spherical, 8-mm radius, masks centered on peak of activity as described above for the model-based fMRI data analysis, limited to the left hemisphere (i.e. AIC: X = −33, Y = 20, Z = −4; dACC: X = −8, Y = 23, Z = 32; VS: X = −12, Y = 11, Z = −8). A further node of the network was also added to include the visual cortex (VC) as a network input, with the ROI centered on coordinates (X = −24, Y = −76; Z = 26), again with 8-mm radius and spherical shape. The principal eigenvariates of these four ROIs were used to extract fMRI time series.

Eight competing models (**Fig. 3**) were defined to be used for family-wise comparisons using a Bayesian random effect approach to determine the causal relations among the three selected ROIs in uncertainty computations. We assumed a single input region in the VC and a fully connected network for the A-matrix (all nodes connected in both directions with all remaining nodes, plus self-connectivity), across all tested models. Eight different architectures were used for the B-matrix, determining differences in the connections targeted by the modulatory signal of the estimated Bayesian uncertainty. The modulatory signal was assumed to always target all the six connections - inwards and outwards - characterizing the VC, plus three connections (single direction) among the remaining three nodes in the network. The resulting eight configurations for the B-Matrix represented all possible target combinations (2 possible directions per 3 pairs of nodes: AIC-dACC, AIC-VS, dACC-VS), allowing grouping the models in families of four, each characterized by one common directed connectivity. Finally, these families were used in three pair-wise comparisons, to test which direction of the modulated connectivity would be characterized by the highest exceedance probability. For instance, the family including the four models characterized by an AIC-to-dACC connectivity was compared with the family characterized by the dACC-to-AIC connectivity.

**Fig. 3.**
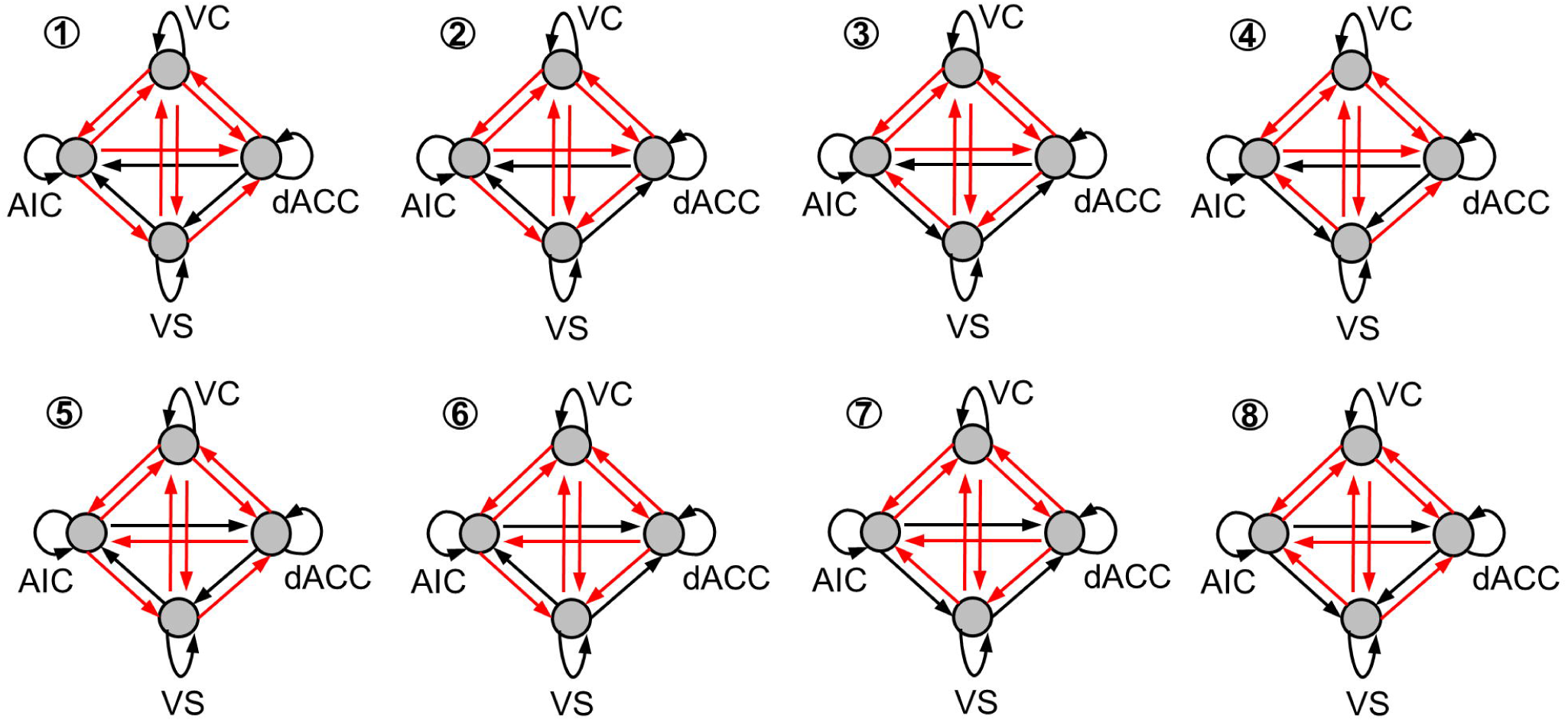
Dynamic causal modeling (DCM) model space. The eight network architectures illustrate the tested DCM models, assuming a fully connected network for all models (illustrated by the presence of black and red arrows) and different connectivity targets for the modulatory signal of model-estimated uncertainty (red arrows). The eight structures illustrate different combinations of these targets of the modulatory signal, to allow for the pair-wise investigation of the directed connectivity among AIC, dACC, and VS. These eight models were grouped into different configuration of four models characterized by a single common directed connectivity, to allow for family-wise comparison. For example, models 1-4 and models 5-8 are respectively characterized by a modulated AIC-to-dACC or dACC-to-AIC connectivity, and varying directed connectivity involving the VS. A comparison between the former and the latter family would then indicate the complementary exceedance probability for AIC-to-dACC and dACC-to-AIC directed connectivity. Similar analyses have been run for the remaining pairs of regions of interest. AIC: anterior insular cortex; dACC: dorsal anterior cingulate cortex, VC: visual cortex; VS: ventral striatum.

### 2.5. Statistical methods

Given the unbalanced sample size, the group difference between HCs and CUD participants was examined by a one-tailed two-sample Welch’s *t*-test. Next, a correlation analysis was conducted to examine how neural activation and connectivity measures correlated with relapse time in CUD patients. To compute the exact *p* value associated with the correlation of coefficient (*r*), we used a non-parametric permutation test (Gu *et al*., 2012; Gu *et al*., 2015). In the permutation test, the null distribution of *r* was first obtained from 10,000 iterations of the permuted data and then used to compute the *p* value associated with the observed *r*. The *α* level was set at *p* = .05.

## 3. Results

### 3.1. Model-agnostic results

We followed previous studies (Rubia *et al*., 2003) in measuring for each group: 1) the mean length of SSD; 2) the mean RTs to go trials and stop failures; and 3) the mean SSRT for both groups, after subtracting the mean length of SSD from the mean RT to the “go trials”. No significant between-group differences were found in the mean length of SSD (Welch’s *t*_40_ = 0.56, *p* = .58), in RTs to the go trials (Welch’s *t*_47_ = 0.18, *p* = .86) or in SSRTs between the two groups (Welch’s *t*_49_ = 0.003, *p* = .997). In conclusion, we found no observable group difference in conventional analyses of the stop signal task.

### 3.2. Uncertainty-related neural activations predict cocaine relapse

Consistent with the literature (Payzan-LeNestour *et al*., 2013; Ma & Jazayeri, 2014; Morriss *et al*., 2019), we found significant activation in the AIC, dACC (extending into the supplementary motor area), and VS that encoded our model-estimated choice uncertainty (**Fig. 2a; Table 2**) at *p*_FWE_ < .05. Notably, the activity of the VS, though less frequently found in association with uncertainty (Zandbelt & Vink, 2010; Ersche *et al*., 2011; Mitchell *et al*., 2014), is well studied in association with SUD and CUD in particular, as a result of over-activation of mesolimbic dopamine signals (Everitt & Robbins, 2016; Koob & Volkow, 2016). In addition, ROI analysis revealed that uncertainty-related activity positively correlated with participants’ days to relapse in left AIC (**Fig. 2b**; *r* = .276, *p* = .0182), dACC (**Fig. 2c**; *r* = .281, *p* = .0187) and bilateral VS (**Fig. 2d-e**; left: *r* = .301, *p* = .0139; right: *r* = .305, *p* = .0125). The group comparison between CUD participants and HCs revealed a significant difference in the right VS (Welch’s *t*_68_ = 2.46, *p* = .02), but not in bilateral AIC (left: Welch’s *t*_68_ = 1.96, *p* = .05; right: Welch’s *t*_60_ = 1.54, *p* = .13), dACC (Welch’s *t*_52_ = 1.92, *p* = .06), or left VS (left: Welch’s *t*_60_ = 0.65, *p* = .52).

**Table 2.**
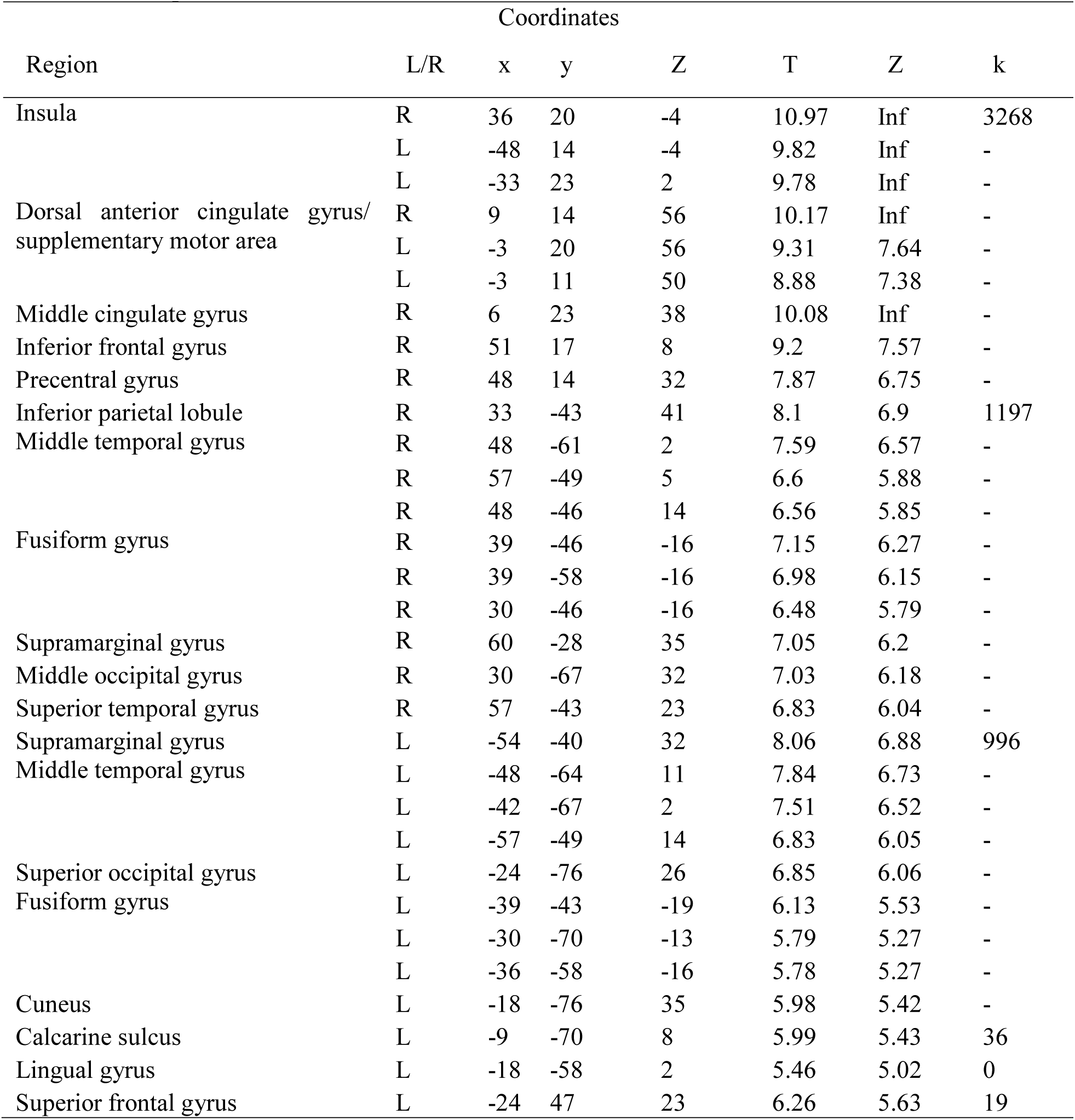
Peaks of regions related to uncertainty in the stop signal task across all participants (*p* < .05 with FWE correction)

| Region | L/R | Coordinates |  |  | T | Z | k |
| --- | --- | --- | --- | --- | --- | --- | --- |
|  |  | x | y | Z |  |  |  |
| Insula | R | 36 | 20 | -4 | 10.97 | Inf | 3268 |
|  | L | -48 | 14 | -4 | 9.82 | Inf | - |
|  | L | -33 | 23 | 2 | 9.78 | Inf | - |
| Dorsal anterior cingulate gyrus/<br>supplementary motor area | R | 9 | 14 | 56 | 10.17 | Inf | - |
|  | L | -3 | 20 | 56 | 9.31 | 7.64 | - |
|  | L | -3 | 11 | 50 | 8.88 | 7.38 | - |
| Middle cingulate gyrus | R | 6 | 23 | 38 | 10.08 | Inf | - |
| Inferior frontal gyrus | R | 51 | 17 | 8 | 9.2 | 7.57 | - |
| Precentral gyrus | R | 48 | 14 | 32 | 7.87 | 6.75 | - |
| Inferior parietal lobule | R | 33 | -43 | 41 | 8.1 | 6.9 | 1197 |
| Middle temporal gyrus | R | 48 | -61 | 2 | 7.59 | 6.57 | - |
|  | R | 57 | -49 | 5 | 6.6 | 5.88 | - |
|  | R | 48 | -46 | 14 | 6.56 | 5.85 | - |
| Fusiform gyrus | R | 39 | -46 | -16 | 7.15 | 6.27 | - |
|  | R | 39 | -58 | -16 | 6.98 | 6.15 | - |
|  | R | 30 | -46 | -16 | 6.48 | 5.79 | - |
| Supramarginal gyrus | R | 60 | -28 | 35 | 7.05 | 6.2 | - |
| Middle occipital gyrus | R | 30 | -67 | 32 | 7.03 | 6.18 | - |
| Superior temporal gyrus | R | 57 | -43 | 23 | 6.83 | 6.04 | - |
| Supramarginal gyrus | L | -54 | -40 | 32 | 8.06 | 6.88 | 996 |
| Middle temporal gyrus | L | -48 | -64 | 11 | 7.84 | 6.73 | - |
|  | L | -42 | -67 | 2 | 7.51 | 6.52 | - |
|  | L | -57 | -49 | 14 | 6.83 | 6.05 | - |
| Superior occipital gyrus | L | -24 | -76 | 26 | 6.85 | 6.06 | - |
| Fusiform gyrus | L | -39 | -43 | -19 | 6.13 | 5.53 | - |
|  | L | -30 | -70 | -13 | 5.79 | 5.27 | - |
|  | L | -36 | -58 | -16 | 5.78 | 5.27 | - |
| Cuneus | L | -18 | -76 | 35 | 5.98 | 5.42 | - |
| Calcarine sulcus | L | -9 | -70 | 8 | 5.99 | 5.43 | 36 |
| Lingual gyrus | L | -18 | -58 | 2 | 5.46 | 5.02 | 0 |
| Superior frontal gyrus | L | -24 | 47 | 23 | 6.26 | 5.63 | 19 |

The leave-one-out SVM classifier showed that it was possible to use the combined ROI results (i.e., left AIC, dACC, and bilateral VS) to predict whether a CUD participant would relapse within 30 days (i.e., early- vs. late-relapse classification, as by definition of early-remission in DSM-IV) with a 78.18% accuracy (**Fig. 2f**). However, we only achieved a close- to-chance level accuracy when applying the same algorithm to classify CUD vs. HC (62.5%).

### 3.3. AIC-to-dACC connection protects against relapse risk

Next, we examined our central question of whether uncertainty-modulated AIC effective connectivity might be associated with relapse. Our family-wise DCM analysis revealed that the family of models where AIC drove dACC activity during uncertainty encoding had an exceedance probability of 98.63% in HCs (**Fig. 4a**). Similar family-wise analysis was used to estimate the exceedance probability associated with the remaining pairs of ROIs, revealing that uncertainty was more likely to modulate AIC-to-VS (90.55%) and VS-to-dACC (66.45%) directed connections in HCs (**Fig. 4a**).

**Fig. 4.**
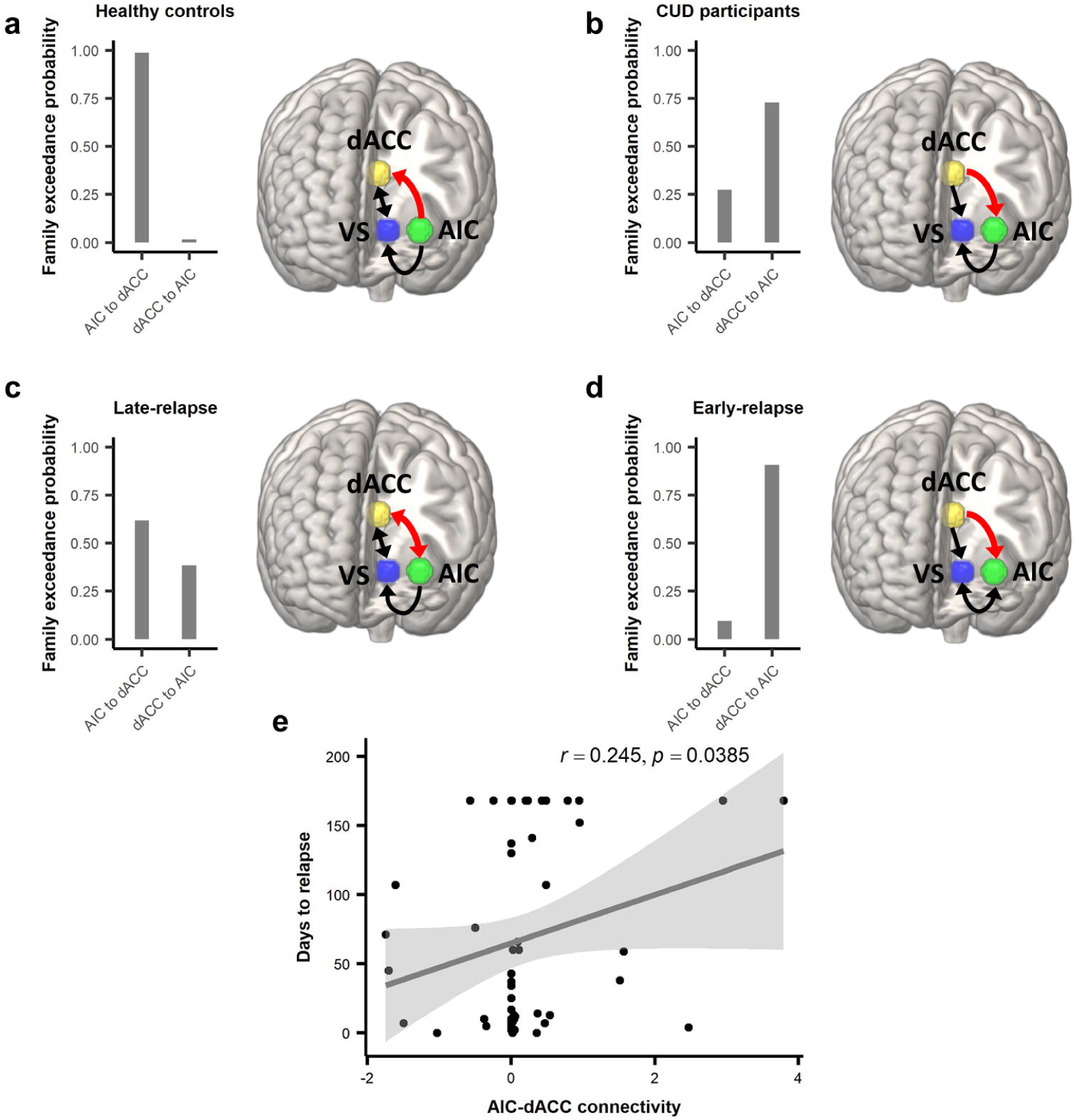
DCM family-wise comparison results. (**a**) The family-wise DCM analysis showed that Bayesian uncertainty is more likely to modulate AIC-to-dACC directed connectivity (98.63% exceedance probability) in healthy controls. (**b**) In contrast, CUD participants showed a dominant dACC-to-AIC directed connectivity (72.76%). This reversal is not present in (**c**) the late-relapse group, which show a mixed result in terms of AIC-dACC connectivity, and it is driven by (**d**) the early-relapse group (with a dACC-to-AIC exceedance probability of 90.56%). Finally, panel (**e**) illustrates that stronger positive uncertainty-modulated AIC-to-dACC connectivity is related to longer abstinence of cocaine (*r* = .245, *p* = .0385).

In the CUD group, in sharp contrast, family-wise comparisons revealed that the signal of uncertainty primarily modulated the directed connectivity from dACC to AIC (72.76%; **Fig. 4b**). We also identified higher exceedance probabilities of models with AIC-to-VS (80.33%) and dACC-to-VS (92.28%) directed connectivity (**Fig. 4b**). To further identify which subgroup drove such pattern, we repeated our family-wise model comparisons after separating CUD early relapsers (<30 days) and CUD individuals with longer abstinence periods, as defined a priori (Adinoff *et al*., 2015; McHugh *et al*., 2016). This analysis showed that the reported reversed connection between AIC and dACC was primarily driven by the early-relapse group (dACC-to-AIC: 90.56%; **Fig. 4d)**, in comparison with those who maintained abstinence beyond 30 days (dACC-to-AIC: 38.34%; **Fig. 4c**). Furthermore, the analysis of the two CUD subgroups also revealed differences in AIC-to-VS (early: 36.731%, late: 93.34%) and the dACC-to-VS connectivity (early: 97.91%, late: 47.49%). Notably, in both connectivity analyses the early relapse group showed the most distant estimates (of the two CUD groups), from HCs estimated connectivity.

These findings highlight a strong link between relapse and the inversion of directed connectivity between AIC and dACC during uncertainty encoding. They also suggest a putative protective role of the AIC-to-dACC connectivity during uncertainty encoding; if this is the case, we speculated we would observe longer abstinence time in those who indeed showed a stronger ACI-to-dACC connection. Therefore, we further tested the relationship between relapse time and subject-specific values estimated for the AIC-to-dACC modulated connectivity. This final analysis confirmed that CUD individuals with stronger positive AIC-to-dACC connection weights remained abstinent for longer periods (*r* = .245, *p* = .0385; **Fig. 4e**). Taken together, these results suggest a protective function of the directional influence from AIC to dACC in cocaine relapse under uncertainty.

## 4. Discussion

Drug addiction is marked by the repeated selection of suboptimal choices despite their adverse consequences (Volkow & Morales, 2015; Everitt & Robbins, 2016). A recent computational model suggests that one of the reasons that can lead an individual to systematically select these suboptimal policies is one’s inability to correctly generate an internal model of the environment (Ognibene *et al*., 2019). This theory postulates that addiction can emerge when environment complexity exceeds, either permanently or temporarily, the cognitive capabilities of an agent, leading to faulty estimations of action-outcome probabilistic associations. In terms of relapse after treatment, this theory predicts that the more these representations are impoverished, the higher the likelihood to reinstate the addictive behavior. Consistent with this prediction, our empirical findings revealed a direct relationship between the neural dynamics of an AIC-driven, uncertainty-computing circuitry and the duration of abstinence post treatment. Specifically, we found that early return to drug use in CUD individuals was predicted by reduced uncertainty-related responses in the AIC and its interconnected regions (dACC and VS). Furthermore, while healthy volunteers showed a predominantly AIC-to-dACC directed connectivity during uncertainty encoding, CUD individuals were characterized by the inverse dACC-to-AIC influential relation, which was mostly driven by early relapsers. Among CUD individuals, those who did have stronger AIC-to-dACC excitatory connections were more likely to remain abstinent for longer periods of time. Collectively, these findings pinpoint the AIC as a key node in driving an uncertainty-encoding neurocircuitry and protecting against relapse.

The insula, in particular the anterior insula, has been under the spotlight in addiction research (Garavan, 2010; Naqvi *et al*., 2014; Koob & Volkow, 2016) primarily due to its role in interoception (Wang *et al*., 2019; Livneh *et al*., 2020) and craving (Garavan, 2010). In a seminal study by Naqvi and colleagues (Naqvi *et al*., 2007), patients with insula damage showed reduced craving for smoking and greater likelihood for quitting, compared to those with lesions outside of the insula. Despite conflicting results from one other human lesion study (Bienkowski *et al*., 2010), the majority of animal models (Contreras *et al*., 2007; Forget *et al*., 2010) and human neuroimaging studies on this topic have independently supported a role of the insula in craving (Filbey *et al*., 2009; Gu *et al*., 2016). In these studies, insula activity is often considered ‘unhealthy’. For example, drug cues induce stronger neural responses in the insula than neutral cues; and consequently, by disrupting the insula, craving reduces and so do drug-seeking behaviors (Contreras *et al*., 2007; Naqvi *et al*., 2007; Forget *et al*., 2010). Different from these previous findings, our results suggest that stronger insula activity and directional influence towards the dACC are ‘healthy’ in the sense that they promote abstinence and protect against relapse.

We reconcile these seemingly disparate findings by reexamining another important line of work on the insula, centering on its role in cognition and decision-making. As introduced earlier, the insula has been consistently involved in the computation of risk, uncertainty, and conflict detection (Clark *et al*., 2008; Preuschoff *et al*., 2008; Bossaerts, 2010; Limongi *et al*., 2013). The central computational element modeled in our study is an individual’s estimation of decision uncertainty, or how likely it is that a choice will lead to any outcome but the desired outcome. This computational process is crucial for optimal decision-making (Fiorillo *et al*., 2003; Niv *et al*., 2005; Platt & Huettel, 2008; Rushworth & Behrens, 2008; Singer *et al*., 2009) and it is distinct from craving, which represents a subjective interoceptive state. Thus, neural responses in the insula might not be simply deemed ‘healthy’ or ‘unhealthy’; instead, the fitness and meaning of insula activity might crucially depend on the task context (e.g. uncertain choices or craving). When drug cues are present, too much insula activity could be detrimental as they signal interoceptive changes related to craving. But when the environment involves no drug stimuli, as is the case in the stop signal task (or in real life when CUD individuals just exited a treatment program), being able to monitor uncertainty requires strong involvement of the insula. In these situations, a dysfunctional AIC-neurocircuitry would then lead to mistakes in uncertainty estimations and therefore increased probability to select actions that are very likely to be associated with ‘unhealthy’ outcomes (Ognibene *et al*., 2019), such as deciding to return to drug-related environments and eventually, relapse.

## Conclusions

Our findings suggest that an insula-driven neurocircuitry that computes decision uncertainty can be an important mechanism and biomarker for CUD early relapse detection and intervention. These results could inform potential early detection of relapse (e.g. by measuring this circuitry during treatment) and intervention (e.g. by giving more intensive treatment and care to individuals with this vulnerability). Thus, these findings could have far-reaching implications for a wide range of addictive disorders beyond CUD, which need to be tested in future studies.

## Acknowledgements

We are grateful to Rani Varghese for her skilled assistance in magnetic resonance imaging scanning and the assistance of the staff on the Substance Abuse Team at the Veterans Affairs North Texas Health Care System, Homeward Bound Inc., and Nexus Recovery Center for their support in the screening and recruitment of study participants.

## Author Contributions

J-CY and VGF analyzed fMRI and behavioral data and wrote the manuscript; RWB and JB collected the data; KR and BA conceived the study, collected the data and edited the manuscript; XG supervised data analysis and wrote the manuscript.

## Data availability statement

Single subject GLM results and associated DCM estimations will be made available upon acceptance of the manuscript in the public repository http://neurovault.org (collection number: --------).

